# Confidence-Guided Machine Learning Staging of the Mouse Estrous Cycle from External Anatomical Images Reduces Cytology Burden

**DOI:** 10.64898/2026.11.21.768307

**Authors:** Zena Del Mundo, Pooya Khosravi, Jocelyn Ha, Antonia Zhang, Avari Alcaire, Abhishek Chanda, Aleksandra E. Sikora, Dequina Nicholas

## Abstract

Although the 2015 National Institutes of Health mandate to consider sex as a biological variable in preclinical studies has increased sex-inclusive study designs, many research groups continue to cite the complexity of monitoring female-specific physiology as a barrier to adoption. Central to this challenge is the murine estrous cycle, a hormonally driven process comprising four stages (proestrus, estrus, metestrus, and diestrus), each associated with distinct hormonal and cytological profiles. Current staging methods rely primarily on vaginal cytology, which requires skilled animal handling, specialized expertise in cytology observation, and considerable time, limiting their routine application outside of reproductive endocrinology. To address these barriers, we developed and evaluated a machine learning framework for estrous cycle staging based on photographs of the mouse external vaginal opening. Using 2,297 photographs from C57BL/6J and BALB/c female mice paired with concurrent cytological assessments and partitioned strictly at the animal level, we trained and evaluated three models of varying granularity: a four-stage model, a three-stage model combining metestrus and diestrus, and a two-stage binary model separating the follicular from the luteal and transitional phases. In a held-out evaluation cohort of 466 photographs from 29 unseen mice, overall classification accuracy was 47.4% (95% CI: 40.7–51.9%) for the four-stage model, 54.5% (47.6–62.9%) for the three-group model, and 69.5% (61.8–75.3%) for the binary two-group model. Misclassifications occurred predominantly between biologically adjacent stages, particularly between proestrus and estrus. Crucially, applying a calibrated confidence threshold to the two-group classifier increased accuracy to 82.8% (95% CI: 71.1–92.1%) among the 43.8% of photographs with confidence ≥ 0.80, and to 92.5% (95% CI: 87.1–96.6%) among the 20.0% of photographs with confidence ≥ 0.90. While external photography does not support standalone four-stage classification, confidence-guided photographic assessment offers an accessible decision-support framework to selectively triage uncertain cases for cytology and reduce the wet-lab burden of estrous monitoring in preclinical research, including cohort assembly for hormone-dependent models of genital tract infection

## Introduction

Rodents are among the most widely used model organisms in biomedical research due to their genetic and physiological similarities to humans. However, the translatability of rodent studies has historically been compromised by the underrepresentation of biological females in experimental designs, particularly in neuroscience and pharmacology [1,2]. Concerns about hormonal variability, higher experimental costs, and a lack of accessible tools to monitor female-specific physiology largely drove the exclusion of female animals. In 2015, the National Institutes of Health (NIH) mandated the consideration of sex as a biological variable in preclinical research, increasing the prevalence of sex-inclusive study designs. The rationale extends beyond neuroscience and pharmacology as sex and the underlying hormonal milieu shape innate and adaptive immune function, susceptibility to infection, and the magnitude and durability of vaccine responses [3]. Nevertheless, adoption has remained uneven, with many research groups citing the added complexity of managing hormonally variable female cohorts as a persistent barrier [4–7].

Even in studies that do include female animals, the estrous cycle is frequently overlooked as a meaningful source of physiological variability. The mouse estrous cycle is a hormonally driven process comprising four discrete stages-proestrus, estrus, metestrus, and diestrus-each defined by a characteristic hormonal profile and cytological composition [8]. Accurate staging is critical across many experimental contexts. In endocrinology and neurobiology, cyclic fluctuations in estrogen and progesterone modulate hormone-secreting cell activity and downstream organ function, whereas in reproductive toxicology, cycling regularity serves as a key endpoint assessment [9]. Failure to account for estrous stage, where relevant, can introduce uncontrolled biological variability that complicates data interpretation and undermines experimental reproducibility.

Importantly, the required staging resolution depends on the experimental workflow. Timed breeding protocols require precise identification of the estrus stage, which represents peak sexual receptivity, or the preceding proestrus stage, during which hormonal priming and reproductive tract remodeling occur. Conversely, experiments requiring hormonal stability, such as drug experiments, tissue collections, and baseline physiological measurements, frequently target the metestrus-to-diestrus window, when circulating ovarian hormone levels are relatively low and stable. For many behavioral and physiological experiments that do not require single-stage precision, broader phase-level classification that separates the follicular phase (proestrus and estrus) from the luteal and transition phase (metestrus and diestrus) may be sufficient [6].

Staging is operationally binding across different mouse models of infectious diseases, particularly female genital tract infection. Rodent models of sexually transmitted infection depend on the hormonal state of the reproductive tract for susceptibility, bacterial burden, and immune readout, and most protocols therefore require cytological screening before an animal can be enrolled. In the widely used β-estradiol-treated mouse model of *Neisseria gonorrhoeae* lower genital tract infection, animals are staged by vaginal cytology, and only those in diestrus are advanced to β-estradiol treatment and inoculation [10–12]. Comparable hormone dependence shapes murine and rat models of *Chlamydia* genital infection [13–16], herpes simplex virus type 2 vaginal challenge [17,18], and colonization by group B *Streptococcus*, *Candida albicans* and *Trichomonas vaginalis* [19–22]*. Neisseria gonorrhoeae* is a WHO high-priority antimicrobial-resistant pathogen, and the mouse vaginal model is the primary *in vivo* platform for testing candidate vaccines and therapeutics [23,24]. However, the throughput cost of daily cytology on large mouse cohorts directly constrains this pipeline.

Because the required staging granularity varies considerably by experimental question, evaluating classification tools across multiple stage groupings is essential.

Current staging methods are technically demanding and represent a practical bottleneck to routine adoption. Vaginal cytology, the gold-standard method, requires skilled animal restraint, precise vaginal lavage or swabbing, slide staining, and accurate microscopic interpretation of mixed cellular populations of nucleated epithelial cells, cornified epithelial cells, and leukocytes [25–28]. Recent machine learning studies, including those by Straus et al. and Sano et al., have demonstrated that deep learning classifiers and object-detection networks can automate stage assignment directly from the stained cytology micrographs with accuracy comparable to human examiners [29–32]. However, while these tools accelerate slide reading, they still require invasive specimen collection and slide preparation for every mouse. Visual observation of the vaginal opening, evaluating tissue swelling, coloration, striation, and moisture, can provide a rapid, non-invasive preliminary assessment prior to lavage, but manual visual staging requires extensive training, varies across animals, and is prone to inter-operator variability [25,33]. In models of genital tract infection, the burden of cytological staging adds an assay-specific cost. Staging requires lavage of the vaginal mucosa that is later infected and sampled for bacterial burden. Repeated pre-enrollment lavage is stressful to the animal [34], may disturb the resident vaginal community that modulates colonization [35,36], and can itself provoke an inflammatory response that confounds the cytology [27]. Lavage also flushes the luminal compartment that supplies the mucosal immune endpoints of these models, including local antibodies, cytokines, and leukocytes, whose levels already vary across the estrous cycle [37,38]. Repeated sampling before challenge therefore risks shifting the baseline against which post-infection responses are measured. Once animals are infected, every lavage is a contact event with an infectious agent under biosafety containment. A non-invasive pre-screen that reduces the number of lavages required to assemble a stage-matched cohort therefore has value beyond operator time.

Applying computer vision to photographs of the external vaginal opening addresses this upstream bottleneck by assessing whether external morphology can guide staging decisions before collecting a cytology specimen. Here, we present a machine learning framework for estrous cycle assessment from standardized mobile-device photographs of the mouse vaginal opening, evaluated across 2,297 paired photographs and cytological preparations from 132 C57BL/6J and BALB/c mice. Using strict animal-level data partitioning, we evaluate region-of-interest (ROI) classifiers across four-stage, three-group (combining metestrus and diestrus), and two-group (proestrus plus estrus versus metestrus plus diestrus) targets, alongside spatial perturbation mapping to examine local anatomical sensitivity. Finally, we demonstrate how calibrated prediction confidence can provide a selective decision-support workflow by identifying high-confidence phase predictions directly while referring the uncertain cases to cytological confirmation.

## Materials and Methods

### Study Population and Reference Cytological Staging

We analyzed a dataset of 2,297 external vaginal opening photographs paired with concurrent vaginal cytology stage labels collected from 132 reproductive aged female C57BL/6J and BALB/c mice. Each mouse contributed a median of 5 photographs (IQR, 4–20; range, 1–105). All animal procedures were approved by the Institutional Animal Care and Use Committee (IACUC) at the University of California, Irvine Protocol #AUP-24-068 and by the IACUC at Oregon State University Protocol #2025-0628.

Vaginal lavages were collected consistently between 8:00 and 9:00 AM using endotoxin-free phosphate-buffered saline (PBS). A P10 pipette was used to dispense 10 µL of PBS, which was gently pipetted up and down 3-5 times at the vaginal opening while avoiding direct contact with the vaginal orifice. The collected lavage fluid was spread across a defined area of a glass microscope slide, air-dried, stained with methylene blue for 15 min, and air-dried prior to imaging on a Hamamatsu NanoZoomer S210 digital slide scanner (CS13239-01) or light microscope. Reference estrous stages were assigned by trained examiners based on the relative proportions of epithelial cells and leukocytes according to established criteria [9–11]: proestrus (P) was defined by a predominance of nucleated epithelial cells; estrus (E) by predominantly anucleated cornified epithelial cells; metestrus (M) by a mixture of nucleated epithelial cells, cornified epithelial cells, and leukocytes; and diestrus (D) by a predominance of leukocytes. Micrographs were used solely to establish ground-truth reference labels and were never provided as inputs to the photographic classifiers.

### Photography and Region-of-Interest (ROI) Preparation

External photographs of the vaginal area were captured using mobile cameras (iPhones and iPads) equipped with a camera-mounted ring light to provide uniform illumination. Mice were gently restrained by the tail with body support. To standardize framing and orientation across variable imaging conditions, an oriented bounding box (OBB) was defined around the vaginal opening using anatomical landmarks. Across the dataset, 1,448 photographs used manually annotated oriented bounding boxes, and 849 photographs were localized using a custom YOLO oriented-bounding-box detector retaining the highest-confidence detection. Each image was rotated to align the bounding box vertically and cropped tightly around the region of interest without extrapolating beyond the box boundaries. Cropped ROIs were resized while preserving aspect ratio and reflection-padded to 448×448 pixels, with 256×256-pixel inputs generated via area-based down sampling. All downstream classification models received only these standardized ROI crops (**Fig. 1A**).

**Fig 1.**
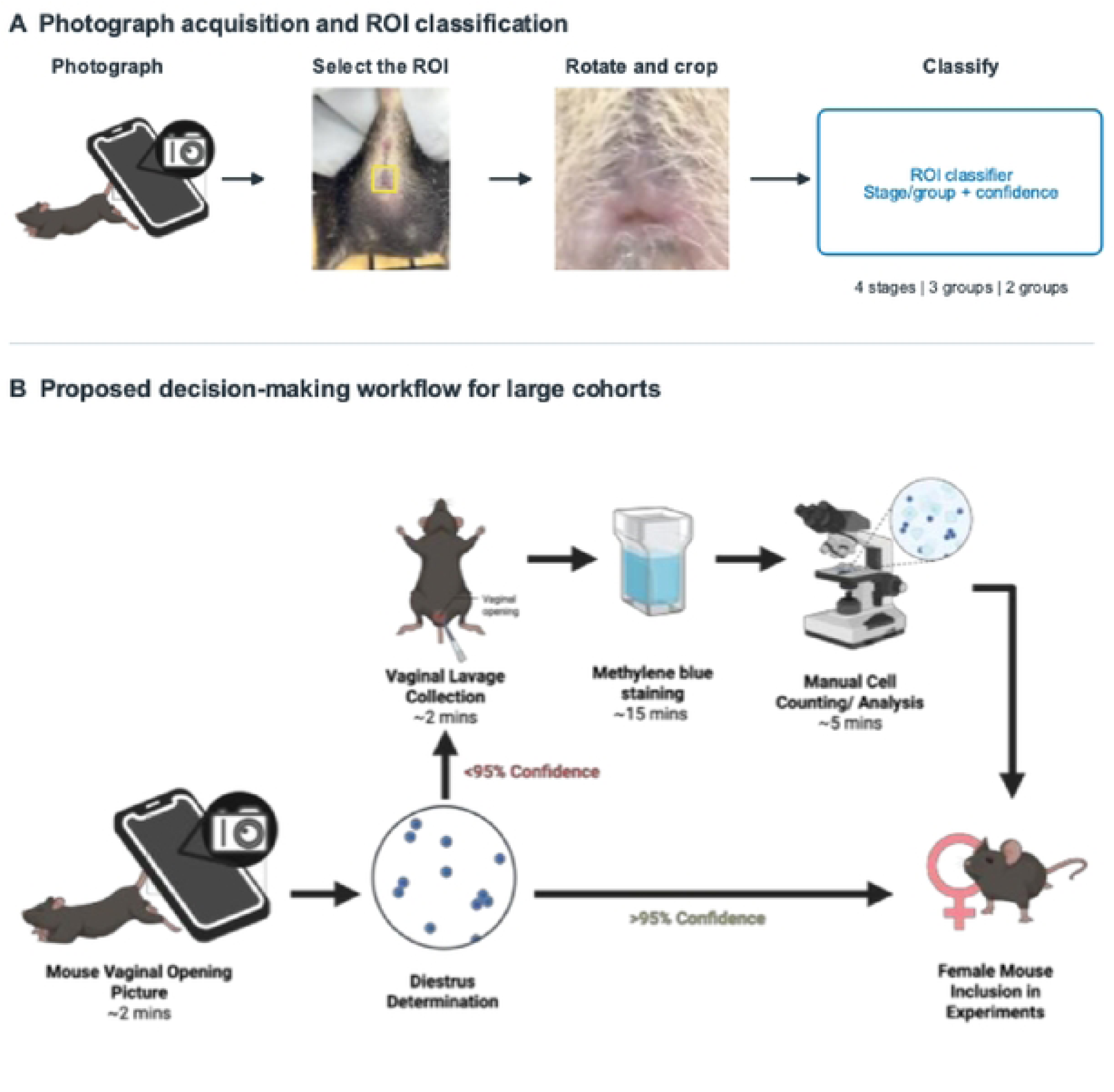
Automated vaginal opening localization and confidence-guided estrous staging workflow. (A) Preprocessing and classification pipeline. Smartphone photographs of the mouse perineum are processed to detect the vaginal opening region of interest (ROI), rotated into a standardized vertical orientation, cropped, and classified at four-stage (proestrus, estrus, metestrus, diestrus), three-group (proestrus, estrus, combined metestrus–diestrus), or binary two-group (proestrus–estrus vs. metestrus–diestrus) resolution. (B) Schematic of the confidence-guided triage workflow for large cohorts. High-confidence photographic predictions (e.g., confidence ≥ 0.80) are accepted directly for experimental assignment, whereas ambiguous, low-confidence cases are triaged to confirmatory vaginal lavage and cytological analysis.

### Dataset Partitioning and Model Development

To prevent animal-level data leakage, all photographs from a given mouse were assigned exclusively to a single data partition. The dataset of 2,297 photographs across 132 mice was divided into a training set of 1,457 photographs (82 mice), a validation set of 374 photographs (21 mice), and a held-out evaluation set of 466 photographs (29 mice). We trained and evaluated classifiers across three staging resolutions derived from the cytological reference labels:

1. Four-stage classification: Proestrus (P), Estrus (E), Metestrus (M), and Diestrus (D) classified independently.
2. Three-group classification: Estrus (E), Proestrus (P), and a combined Metestrus + Diestrus (M+D) group.
3. Two-group (binary phase) classification: Combined Estrus + Proestrus (E+P; follicular phase) versus combined Metestrus + Diestrus (M+D; luteal and transitional phases).

Classification models were built using an EfficientNetV2-S convolutional backbone [39] (alongside a DINOv3 Small vision transformer backbone with spatial attention [40] for four-stage classification) initialized with pretrained weights. Models were fine-tuned on the training partition for up to 24 epochs using the AdamW optimizer, class-weighted cross-entropy loss, and label smoothing, with standard geometric and color augmentations (random horizontal flips, small rotations, and modest color variation). The validation partition was used for early stopping (6 epochs patience), checkpoint selection based on macro-F1, and post-hoc temperature scaling [41] to calibrate predicted class probabilities. Model confidence was defined as the maximum calibrated probability assigned to a predicted class or group. Full architecture, cross-validation model selection, and calibration details are provided in S1 Methods.

### Statistical Analysis and Spatial Sensitivity

Performance on the held-out evaluation set (466 photographs from 29 mice) was assessed using accuracy, balanced accuracy, macro-F1, class-specific sensitivity and precision, one-versus-rest area under the receiver operating characteristic curve (AUC), and average precision (AP)). Non-parametric 95% confidence intervals (CIs) were calculated via 2,000 bootstrap samples clustered at the mouse level (retaining all photographs from each sampled animal). For selective prediction triage, we evaluated accuracy and retained-image coverage (the percentage of photographs meeting or exceeding a given confidence cutoff) across confidence thresholds from 0.0 to 1.0, highlighting illustrative operating cutoffs of ≥ 0.80 and ≥ 0.90. Differences in prediction correctness across cytological stages and collection cohorts were evaluated using mouse-clustered binomial generalized estimating equations (GEE) with Holm multiplicity correction.

To examine whether stage predictions relied on anatomically relevant regions of the vaginal opening, we performed a spatial occlusion sensitivity analysis by replacing one ROI grid tile at a time with Gaussian-blurred pixels (σ = 12 pixels at 256 × 256 resolution, σ = 21 pixels at 448 × 448 resolution) and measuring the resulting decrease in reference-stage probability. Aggregate effects were quantified across all evaluation photographs using a 3 × 3 grid, and fine-grained 7 × 7 sensitivity maps were generated for the representative correctly classified and misclassified examples.

## Results

### Estrous Cycle Staging Dataset and Vaginal Opening Identification

To develop and evaluate an automated estrous cycle staging framework, we curated a dataset of 2,297 vaginal opening photographs paired with concurrent vaginal cytology assessments from 132 reproductive-age female C57BL/6J and BALB/c mice. Each mouse contributed a median of 5 photographs (IQR, 4–20; range, 1–105). Cytology slides were evaluated by trained human examiners to establish reference stage labels, and each photograph was cropped and standardized around the vaginal opening region of interest (ROI) using anatomical landmarks (**Fig 1A and Fig 2**). To prevent animal-level data leakage across splits, all photographs from a given mouse were assigned exclusively to a single partition: 1,457 photographs from 82 mice in the training set, 374 photographs from 21 mice in the validation set, and 466 photographs from 29 unseen mice in the held-out evaluation set (**Table 1**). Within the 466-photograph evaluation set, reference cytological labels comprised 119 diestrus, 177 estrus, 114 metestrus, and 56 proestrus images.

**Fig 2.**
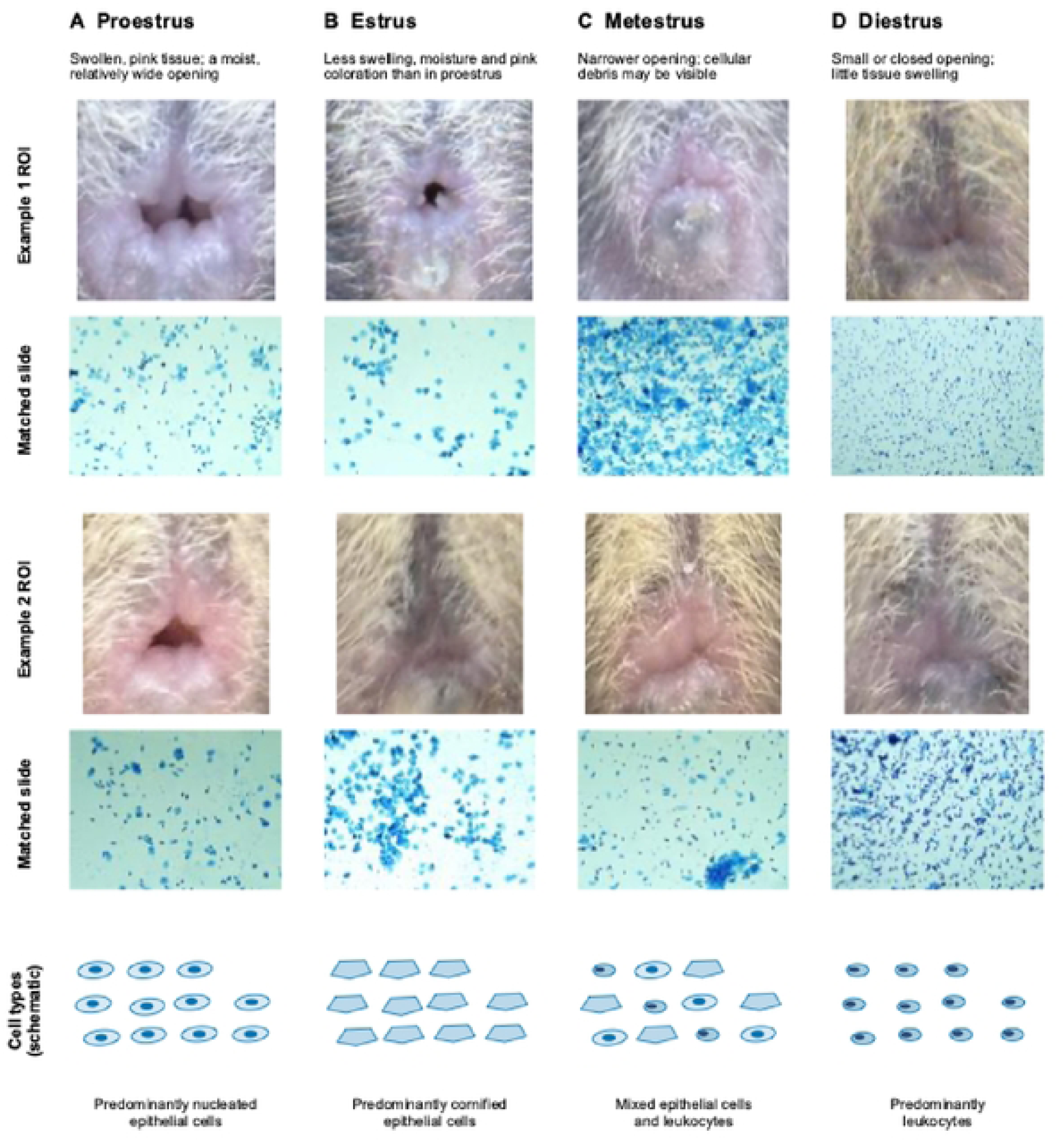
Representative external vaginal opening morphology and paired vaginal cytology across the mouse estrous cycle. Columns depict the four estrous stages: (A) proestrus, characterized externally by swollen, pink, moist labial tissue with a wide vaginal opening and cytologically by predominantly nucleated epithelial cells; (B) estrus, showing reduced swelling and pallor with predominantly anucleated cornified epithelial cells; (C) metestrus, showing a narrower opening with visible cellular debris and a mixed population of epithelial cells and leukocytes; and (D) diestrus, characterized by a small or closed opening with minimal swelling and predominantly leukocytes. Top and middle rows show two independent examples of standardized vaginal opening ROIs paired with concurrent methylene blue–stained vaginal smears from the same animal and sampling session; the bottom row illustrates characteristic cytological cell types.

**Table 1.** Distribution of vaginal opening photographs and unique mice across estrous stages and dataset partitions.

| **Partition** | **Unique Mice** | **Total Photographs** | **Diestrus (D)** | **Estrus (E)** | **Metestrus (M)** | **Proestrus (P)** |
|---|---|---|---|---|---|---|
| Training | 82 | 1,457 | 349 | 574 | 355 | 179 |
| Validation | 21 | 374 | 89 | 152 | 90 | 43 |
| Development Total (Train + Val) | 103 | 1,831 | 438 | 726 | 445 | 222 |
| Evaluation (Held-out) | 29 | 466 | 119 | 177 | 114 | 56 |
| All Partitions Combined | 132 | 2,297 | 557 | 903 | 559 | 278 |

*All photographs from an individual mouse were assigned exclusively to a single partition to prevent animal-level data leakage. Each mouse contributed a median of 5 photographs (IQR, 4–20; range, 1–105).*

### Classification Performance Across Three Staging Resolutions

We evaluated photographic classification across three experimentally relevant target definitions on the held-out evaluation cohort of 466 photographs from 29 mice: a four-stage model classifying proestrus, estrus, metestrus, and diestrus individually; a three-group model combining metestrus and diestrus into a single group while classifying proestrus and estrus independently; and a binary two-group model separating the follicular phase (proestrus plus estrus) from the luteal and transitional phases (metestrus plus diestrus). Across the three staging resolutions, overall classification accuracy increased progressively as stage boundaries were consolidated into broader functional groups, rising from 47.4% (95% CI: 40.7–51.9%) for four stages to 54.5% (95% CI: 47.6–62.9%) for three groups and 69.5% (95% CI: 61.8–75.3%) for two groups (**Table 2**). Balanced accuracy similarly increased from 43.3% (95% CI: 37.4– 48.9%) in the four-stage task to 47.5% (95% CI: 41.3–56.1%) in the three-group task and 69.5% (95% CI: 62.2–75.2%) in the two-group task, with corresponding macro-F1 scores of 0.434, 0.473, and 0.695, respectively (**Table 2**; complete metrics in **Table S1** and discrimination curves in **Fig. S1**).

**Table 2.**
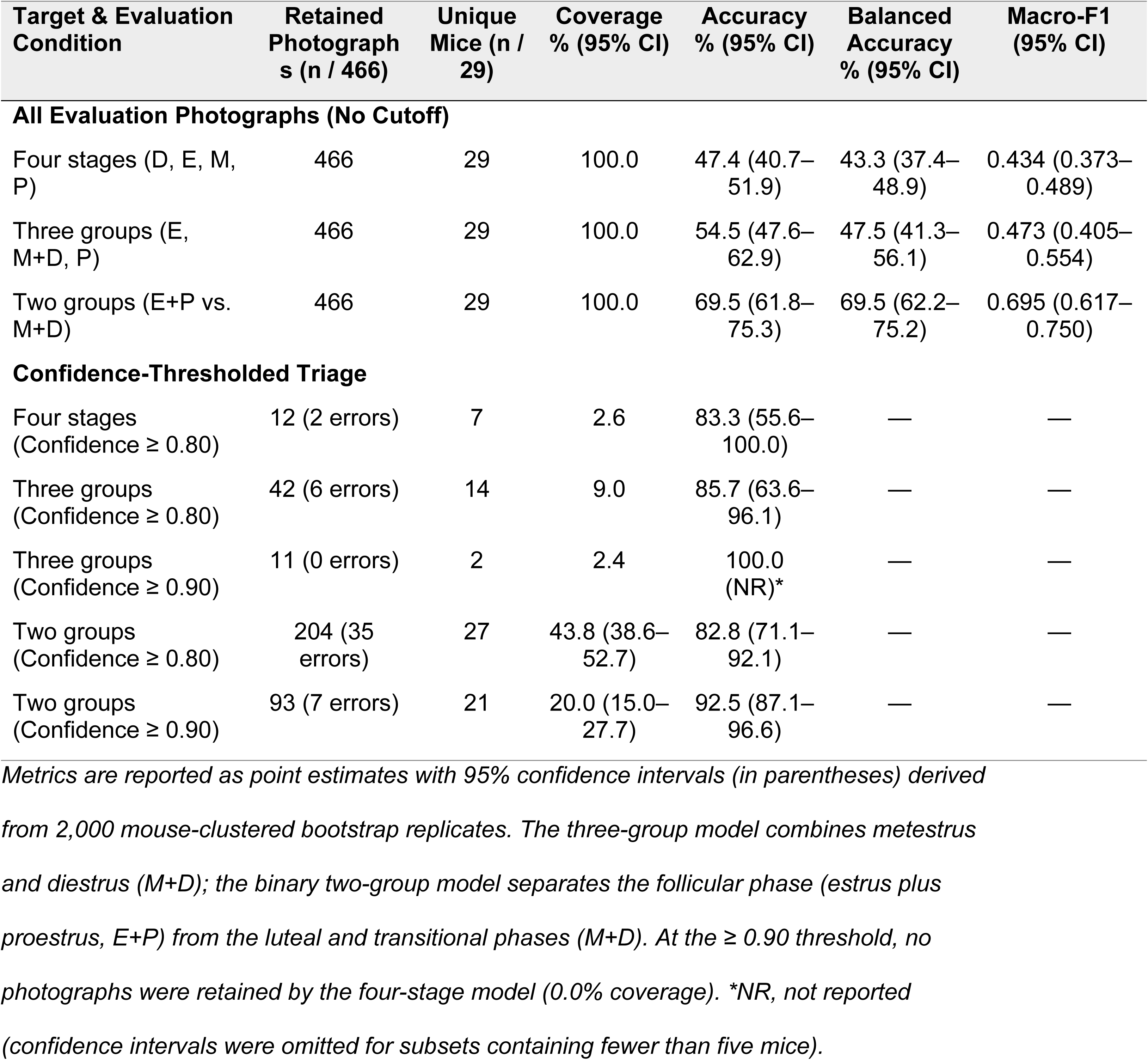
Classification performance and confidence-thresholded triage outcomes on the held-out evaluation cohort (n = 466 photographs from 29 mice).

Inspection of the confusion matrices and class-specific performance metrics illustrated the patterns underlying these accuracy differences (**Fig. 3**). In the four-stage model, class-specific sensitivity was 53.7% (95% CI: 37.1–61.4%; precision: 53.1%) for estrus, 50.0% (95% CI: 35.5–62.3%; precision: 43.5%) for metestrus, and 47.9% (95% CI: 31.2–68.8%; precision: 47.1%) for diestrus, but only 21.4% (95% CI: 13.0–37.0%; precision: 34.3%) for proestrus (**Fig. 3A, D**). Misclassifications occurred predominantly between biologically adjacent stages: most notably, 30 of the 56 proestrus photographs (53.6%) were classified as estrus, compared with 8 (14.3%) classified as metestrus and 6 (10.7%) as diestrus. Because metestrus is a brief transitional stage whose cytological and external morphological profile blends into diestrus, combining metestrus and diestrus in the three-group model substantially improved identification of the combined M+D group, achieving a sensitivity of 64.8% (95% CI: 51.2–86.6%) and precision of 67.4% (95% CI: 56.9–74.4%; AUC: 0.73; average precision [AP]: 0.72), whereas sensitivity remained 49.2% (precision: 48.3%) for estrus and 28.6% (precision: 25.8%) for proestrus (**Fig. 3B, E and Fig. S1**). Grouping proestrus with estrus in the binary two-group model resolved the frequent proestrus–estrus boundary confusion and yielded balanced performance across both functional phases, with a sensitivity of 73.8% (95% CI: 57.0–81.5%; precision: 68.0%; AUC: 0.78; AP: 0.79) for estrus plus proestrus and 65.2% (95% CI: 53.1–81.6%; precision: 71.4%; AUC: 0.78; AP: 0.77) for metestrus plus diestrus (**Fig. 3C, F and Fig. S1**).

**Fig. 3.**
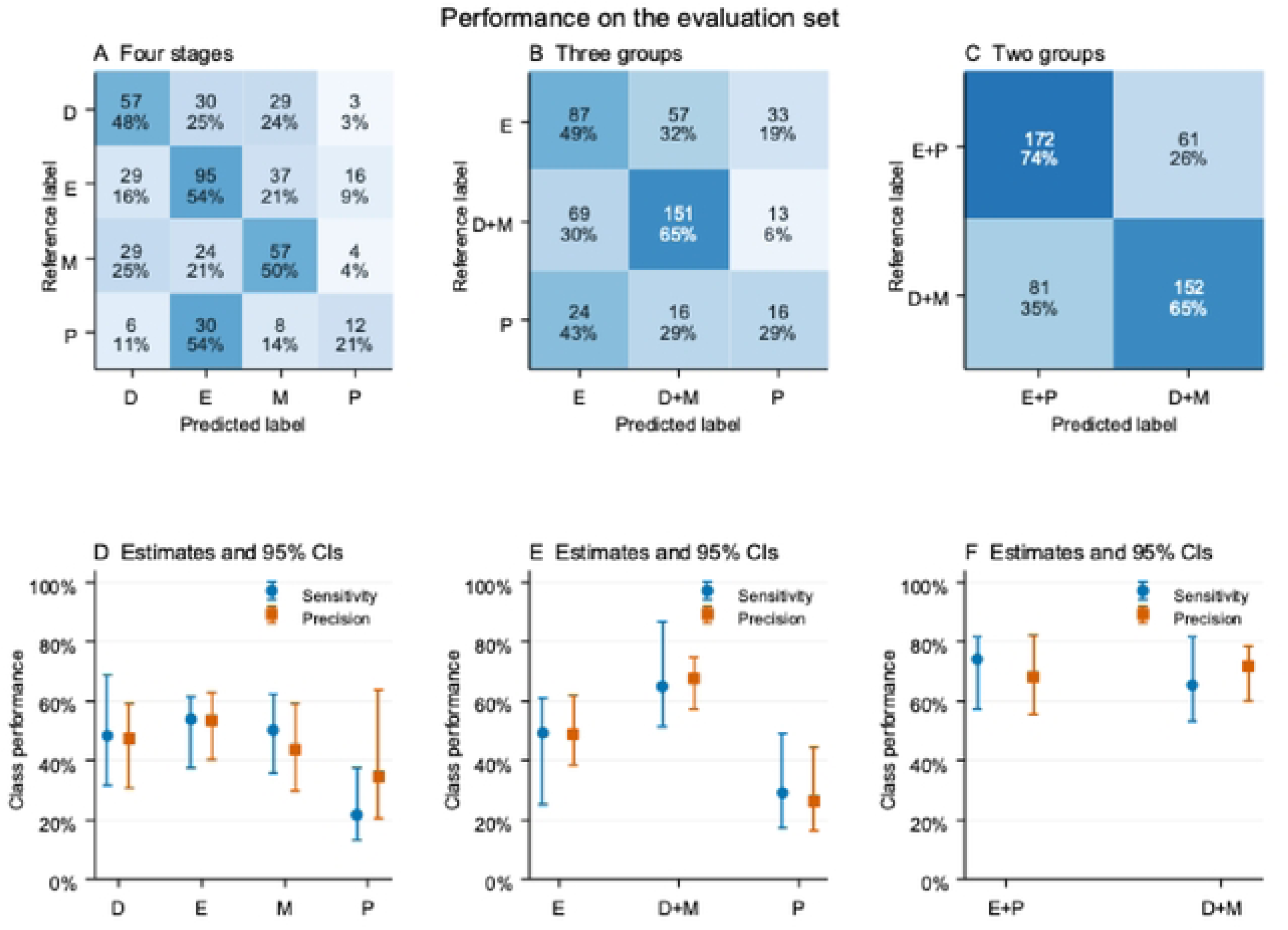
Evaluation set classification performance across four-stage, three-group, and binary two-group staging resolutions. (A–C) Confusion matrices for the held-out evaluation cohort (n = 466 photographs from 29 mice) across the (A) four-stage, (B) three-group, and (C) binary two-group classifiers. Cell values report photograph counts and row-normalized percentages within each cytological reference class. (D–F) Class-specific sensitivity and precision for the (D) four-stage, (E) three-group, and (F) two-group models. Error bars represent 95% confidence intervals estimated from 2,000 mouse-clustered bootstrap replicates. D, diestrus; E, estrus; M, metestrus; P, proestrus.

### Confidence Calibration and Selective Prediction Triage

Because external anatomical transitions between estrous stages are continuous, photographs captured near stage transitions present inherently ambiguous visual features. We therefore evaluated whether calibrated prediction confidence could reliably isolate high-accuracy predictions from uncertain cases requiring cytological confirmation (**Fig. 4**). Temperature scaling fitted on the validation set improved two-group probability calibration on the evaluation cohort— reducing log loss from 0.585 to 0.574, Brier score from 0.399 to 0.393, and expected calibration error (ECE) from 0.066 to 0.057—resulting in close agreement between predicted confidence and observed accuracy across probability bins (**Fig. 4D**).

**Fig 4.**
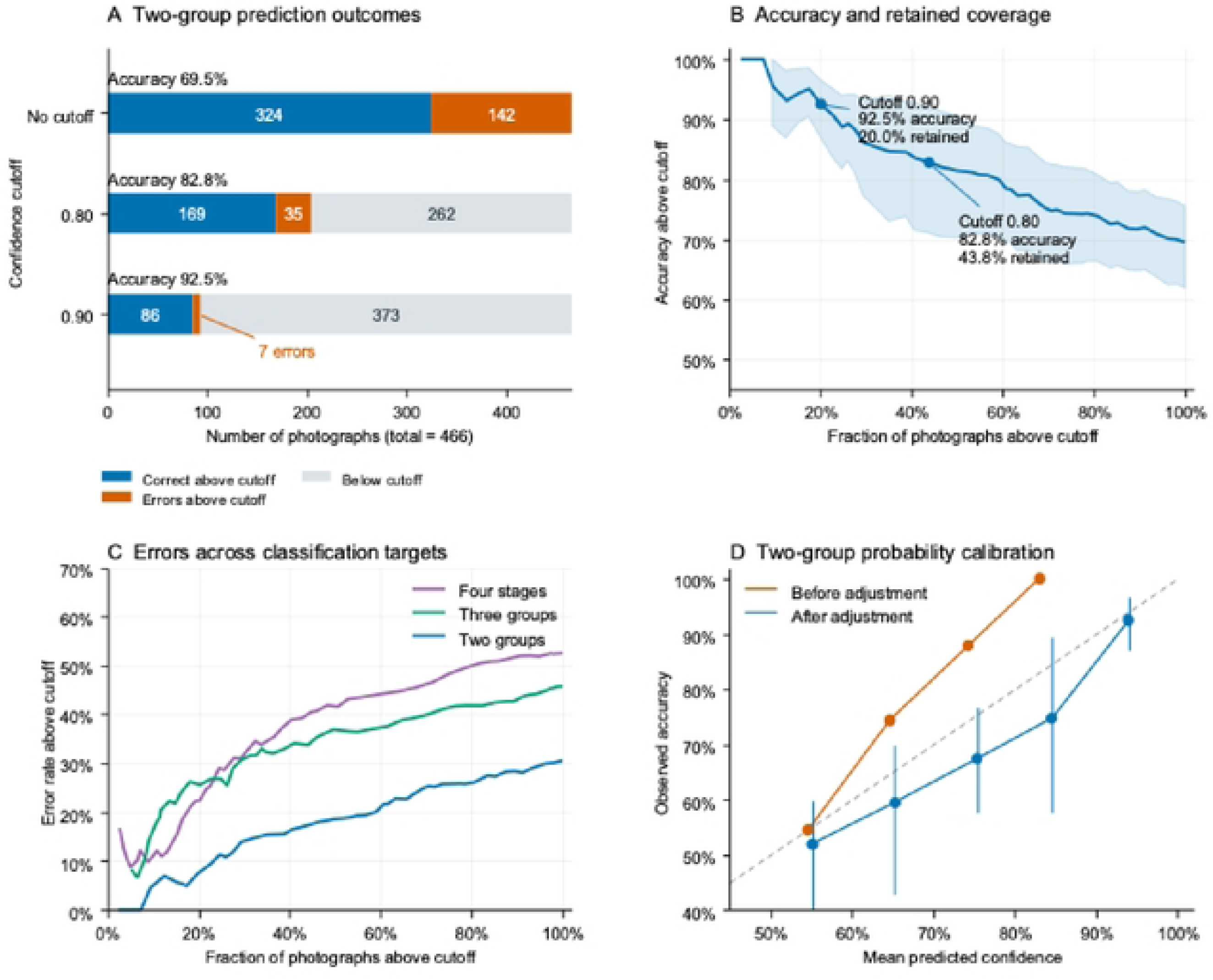
Selective prediction trade-offs and probability calibration for confidence-guided triage. (A) Distribution of evaluation set outcomes for the binary two-group classifier across all photographs (no cutoff; n = 466) and at confidence thresholds of ≥ 0.80 and ≥ 0.90, illustrating correct retained predictions, retained errors, and below-threshold photographs referred for cytology. (B) Classification accuracy among retained photographs as a function of retained-image coverage for the two-group model; shaded bands denote pointwise 95% mouse-bootstrap confidence intervals. (C) Error rate among retained predictions versus coverage across the four-stage, three-group, and two-group classifiers. (D) Reliability diagram for the two-group classifier comparing mean predicted confidence against observed accuracy across probability bins before and after validation-fitted temperature scaling; the dashed diagonal represents ideal calibration.

Applying confidence thresholds to the two-group model produced a strong accuracy–coverage trade-off (**Table 2 and Fig. 4A–C**). Across all 466 evaluation photographs (100% coverage), the unthresholded two-group model yielded 324 correct predictions and 142 errors (69.5% accuracy). Restricting direct classification to photographs with predicted confidence ≥ 0.80 retained 204 photographs (43.8% coverage; 95% CI: 38.6–52.7%) from 27 mice, yielding 169 correct predictions and 35 errors for an accuracy of 82.8% (95% CI: 71.1–92.1%). Increasing the confidence cutoff to ≥ 0.90 retained 93 photographs (20.0% coverage; 95% CI: 15.0–27.7%) from 21 mice, with 86 correct predictions and only 7 errors, corresponding to an accuracy of 92.5% (95% CI: 87.1–96.6%) (**Fig. 4A, B**). By contrast, the four-stage and three-group models produced substantially fewer high-confidence predictions (**Fig. 4C**): at the ≥ 0.80 threshold, the four-stage model retained 12 photographs from 7 mice (2.6% coverage; 83.3% accuracy; 2 errors) and the three-group model retained 42 photographs from 14 mice (9.0% coverage; 85.7% accuracy; 6 errors; and 11 photographs from 2 mice with 0 errors at the ≥ 0.90 cutoff; **Table 2**).

### Stage and Collection Cohort Comparisons

To test whether model performance differed across data collection cohorts or cytological reference stages, we analyzed prediction correctness using mouse-clustered binomial generalized estimating equations (GEE) adjusting additively for stage and cohort (**Fig. S2 and Table. S2**). Prediction accuracy did not differ significantly across the four collection cohorts for any classification resolution (Holm-adjusted omnibus P = 0.394 for four-stage, three-group, and two-group models). Conversely, prediction correctness differed significantly across reference stages after adjusting for collection cohort (Holm-adjusted omnibus P = 0.0129 for four stages, P = 6.62 x 10^-6 for three groups, and P = 0.0129 for two groups), consistent with the lower classification sensitivity observed during proestrus.

### Spatial Sensitivity to Local Anatomical Detail

To verify that classification decisions relied on biologically relevant features of the vaginal opening rather than background artifacts, we quantified local spatial sensitivity by replacing individual grid tiles of each ROI with Gaussian-blurred pixels and measuring the resulting change in reference-stage probability (**Fig. 5 and Fig. S3**). Across all 466 evaluation photographs on a 3 × 3 grid, blurring the lower-center tile produced the largest mean decrease in reference-stage probability for proestrus (0.065; 95% CI: 0.042–0.097; n = 11 mice) and diestrus (0.047; 95% CI: 0.030–0.063; n = 20 mice), whereas blurring the central tile produced the largest mean decrease for metestrus (0.046; 95% CI: 0.031–0.062; n = 20 mice). In estrus photographs, blurring the central tile increased the predicted estrus probability by an average of 0.061 (original minus perturbed probability of-0.061; 95% CI:-0.078 to-0.042; n = 24 mice), indicating that attenuating fine central texture shifted model predictions toward estrus. High-resolution 7 × 7 perturbation maps of representative correctly classified photographs (**Fig. 5**) confirmed that positive sensitivity localized directly over the vaginal orifice and surrounding labial tissue in proestrus, metestrus, and diestrus, whereas incorrectly classified examples (**Fig. S3**) displayed mixed or opposing local probability shifts near the opening.

**Fig. 5.**
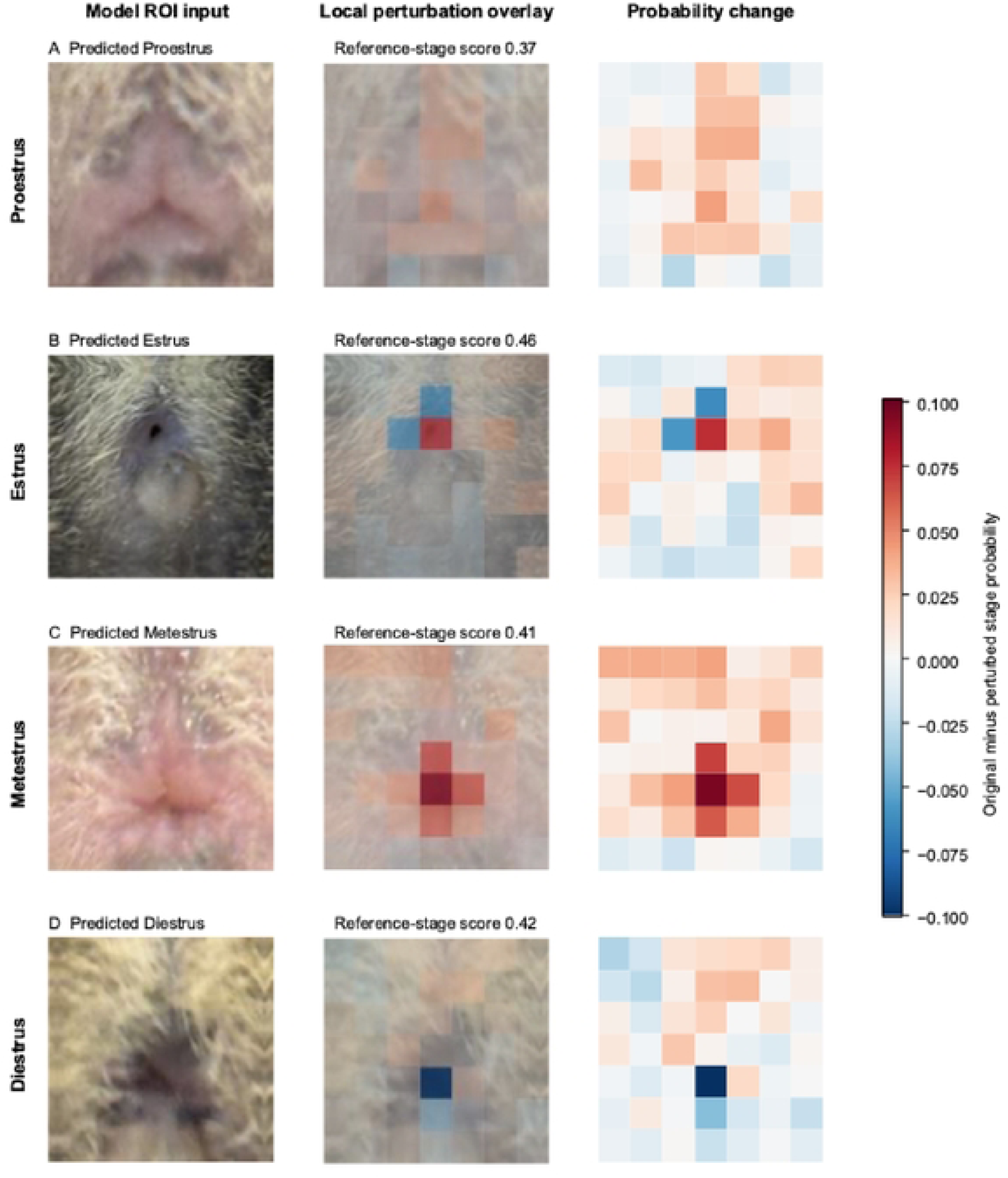
Spatial occlusion sensitivity of the four-stage classifier across correctly staged vaginal opening images. Representative examples of correctly classified photographs in (A) proestrus, (B) estrus, (C) metestrus, and (D) diestrus. For each stage, panels display the standardized ROI input (left), a 7 × 7 local Gaussian-blur perturbation overlay (middle), and the resulting change in predicted probability for the reference stage (right; original minus blurred probability). Warm (red) regions indicate anatomical areas where local detail supports the true stage classification (blurring reduces reference-stage probability), whereas cool (blue) regions indicate areas where blurring increases reference-stage probability.

## Discussion

Among the three classification resolutions evaluated on unseen animals, photographic staging accuracy increased progressively from 47.4% in the four-stage model to 54.5% in the three-group model and 69.5% in the binary two-group model, which separates the follicular phase (proestrus plus estrus) from the luteal and transitional phases (metestrus plus diestrus). This performance progression reflects the biological continuity of the rodent estrous cycle, in which neighboring stages share overlapping morphological and cytological features that create ambiguous boundaries, even for experienced human examiners. Because transitions between stages occur gradually, a single photograph captured near a stage transition may exhibit external tissue characteristics consistent with more than one classification, placing a practical ceiling on standalone four-stage accuracy from external images alone. In our four-stage evaluation, misclassifications were heavily concentrated between biologically consecutive stages—most prominently proestrus photographs classified as estrus (30 of 56 images) and reciprocal errors among estrus, metestrus, and diestrus. Combining metestrus with diestrus in the three-group model, and further uniting proestrus with estrus in the binary model, progressively reduces this inter-stage boundary overlap and confirms that morphological continuity between adjacent stages is the primary driver of misclassification.

Given the limited sensitivity for individual transitional stages such as proestrus, we do not recommend external photography as a standalone replacement for vaginal cytology when four-stage precision is required. However, our confidence-calibration analysis demonstrates that photographic classifiers have substantial value as decision-support triage tools that can selectively reduce the burden of cytology in experiments requiring broad phase grouping. Specifically, when distinguishing the follicular phase (proestrus and estrus) from the luteal and transitional phases (metestrus and diestrus), applying a calibrated confidence threshold of ≥ 0.80 increased classification accuracy from 69.5% to 82.8% across the 43.8% of photographs meeting the cutoff, while a threshold of ≥ 0.90 achieved 92.5% accuracy (with only 7 errors across 93 retained photographs) at 20.0% coverage. In a practical implementation (**Fig. 1B**), each photographic prediction is paired with a calibrated confidence score; investigators can prespecify a confidence cutoff aligned with their experiment’s error tolerance, proceed directly with experimental procedures for animals whose predictions meet or exceed that cutoff, and perform vaginal lavage and microscopic cytology solely for animals with lower-confidence predictions.

This upstream photographic triage approach complements automated deep learning tools developed for vaginal cytology micrographs. Whereas automated cytology classifiers accelerate slide interpretation after a vaginal smear has been collected, stained, and imaged, photographic assessment occurs before specimen collection to determine whether a lavage is needed. Furthermore, our ROI standardization pipeline and spatial perturbation analyses (**Fig. 5**) demonstrate that predictions are driven by local contours and tissue appearance around the vaginal opening, consistent with published visual staging criteria, and perform consistently across independent collection cohorts without detectable batch drift (**Fig. S2**). Standardizing ROI orientation and cropping across mobile devices also facilitates the scalable collection of larger, multi-cohort datasets to support future model refinement.

The confidence-guided workflow maps directly onto cohort assembly for mouse models of sexually transmitted infection. Enrollment in the estradiol-treated gonococcal model requires identifying animals in diestrus, and chlamydial, herpes and fungal vaginal models similarly require a defined hormonal state at challenge [11,12,16,18,21]. In this setting the binary follicular versus luteal and transitional output is the decision of interest, and the asymmetric cost structure favors the selective-prediction approach, as photographs classified with high confidence as metestrus or diestrus can be advanced to lavage-confirmed enrollment, while the remainder are either deferred to the next day or triaged to immediate cytology. The external features the classifier uses, namely swelling, pallor, cornified surface texture and aperture size, are themselves downstream of the estrogen-and progesterone-driven epithelial remodeling that governs mucosal susceptibility [38,42]. Therefore, the photograph reports on the barrier state that determines infection outcome rather than on an arbitrary label.

Stage-aware cohort assembly also addresses an interpretive problem in this literature, where unstaged or incompletely staged cohorts carry hormonally driven variance in bacterial burden and in local antibody and leukocyte responses [15,37]. In the mouse vagina, luminal neutrophil infiltration tracks ovarian cycle phase and is largely independent of pathogen challenge across models of *N. gonorrhoeae*, *Candida albicans*, *Trichomonas vaginalis,* and herpes simplex virus type 2 infection [43], and physiological progesterone surges drive neutrophil influx into the lower genital tract [44]. In an unstaged cohort, cycle phase contributes to the local inflammatory signal attributed to infection, and stage-matched enrollment directly controls for it. The same hormonal dependence accompanies cyclical and contraceptive-driven changes in susceptibility to gonococcal, chlamydial and viral infection in women [3,45–47].

Several limitations of the present study should be noted and help define priorities for future work. First, although assigning all photographs from each mouse to a single partition prevented animal-level leakage and mouse-clustered bootstrapping accounted for repeated photographs per animal, our evaluation partition was inspected during earlier project development. Thus, these findings represent a retrospective evaluation and warrant prospective validation in an independent cohort. Second, model performance was evaluated in two inbred mouse strains, C57BL/6J and BALB/c, and ground-truth reference labels were derived from human cytological assessments without formal multi-rater agreement measurements. Because human cytological staging is itself subject to inter-observer variability during stage transitions, incorporating consensus multi-rater cytology labels and expanding training datasets to additional mouse strains with diverse coat pigmentation and perineal morphology will be important next steps. Finally, because multiple photographs in our dataset could correspond to a single sampling occasion, retaining 20.0% or 43.8% of photographs above a confidence cutoff does not translate one-to-one into the proportion of cytology specimens avoided. A prospective workflow study that captures a single standardized photograph per animal per session and directly measures avoided lavage procedures, and staff time saved will be essential to quantify real-world operational savings. All photographs in this study were collected from uninfected, hormonally untreated animals. Exogenous estradiol, inflammatory discharge, antibiotic co-treatment, and infection-associated tissue changes alter external perineal appearance, so applying this to infection models requires prospective validation in hormone-treated and infected cohorts before using photographic triage after challenge rather than only at enrollment.

Together, our findings establish an accessible, confidence-guided computational framework that can help lower practical barriers to estrous cycle monitoring in sex-inclusive biomedical research.

## Data Availability

The minimal dataset is available at Dryad via https://doi.org/10.5061/dryad.xpnvx0kzj

## Supporting Information Captions

**Fig. S1. Receiver operating characteristic and precision–recall curves across classification resolutions.** One-versus-rest receiver operating characteristic (ROC; top) and precision–recall (PR; bottom) curves on the held-out evaluation cohort (n = 466 photographs from 29 mice) for the four-stage, three-group, and binary two-group models. Insets report per-class area under the ROC curve (AUC) and average precision (AP).

**Fig. S2. Classification accuracy across collection cohorts and cytological reference stages.** Accuracy point estimates and 95% mouse-bootstrap confidence intervals stratified by imaging cohort (top) and cytological reference stage (bottom) for the four-stage, three-group, and two-group classifiers. Holm-adjusted omnibus P values were derived from mouse-clustered binomial generalized estimating equations.

**Fig. S3. Spatial occlusion sensitivity for misclassified vaginal opening images.** Representative 7 × 7 local Gaussian-blur perturbation maps for one misclassified evaluation photograph per reference stage (selected near the median reference-stage probability among incorrect predictions). Panel layout and color scale match Fig 3.

**Table S1. Additional overall and class-specific classification performance.** Values are estimates (95% confidence intervals) for 466 photographs from 29 mice, without a confidence cutoff. Intervals use 2,000 bootstrap samples of mice, retaining all photographs from each sampled mouse. Overall accuracy, balanced accuracy and macro-F1 are reported in Table 2.

Four stages: three EfficientNetV2-S and three DINOv3 spatial classifiers; three groups: one EfficientNetV2-S; two groups: three EfficientNetV2-S classifiers. Ensemble components have equal weights. Probability-based metrics precede final task-level temperature scaling.

Sensitivity is recall; precision is positive predictive value. AUROC, area under the receiver operating characteristic curve. Mean per-mouse accuracy weights mice equally; other metrics weight photographs equally. Brier score is the mean sum of squared class-probability errors. Lower Brier score and negative log likelihood indicate better probability predictions. Confidence intervals do not include model-selection or reference-label uncertainty.

**Table S2. Associations of reference stage and collection cohort with classification correctness.** Analysis included 466 photographs from 29 mice. Binomial generalized estimating equations modeled correct classification using reference stage and collection cohort jointly, with a logit link, independent working correlation and mouse-clustered sandwich covariance. Model procedures are specified in S1 Table.

Odds ratios compare the first group with the second, adjusting for the other factor; values above 1 indicate higher odds of correct classification in the first group. Parentheses contain unadjusted 95% Wald confidence intervals (CIs). Overall tests are Wald tests with 3 degrees of freedom; an overall test has no single odds ratio (—).

All P values are Holm-adjusted: six overall tests form one family and 36 pairwise comparisons form a separate family across the three tasks. Cohorts 1–4 identify image collections, not verified strains or treatment groups. Stage comparisons use the four original cytological stages. These exploratory tests do not establish biological effects or cohort equivalence.

